# DIA-based Proteomics Identifies IDH2 as a Targetable Regulator of Acquired Drug Resistance in Chronic Myeloid Leukemia

**DOI:** 10.1101/2021.04.08.438976

**Authors:** Wei Liu, Yaoting Sun, Weigang Ge, Fangfei Zhang, Lin Gan, Yi Zhu, Tiannan Guo, Kexin Liu

**Author notes:** Correspondence (T.G.); (K.L.).

## Abstract

Drug resistance is a critical obstacle to effective treatment in patients with chronic myeloid leukemia (CML). To understand the underlying resistance mechanisms in response to imatinib (IMA) and adriamycin (ADR), the parental K562 cells were treated with low doses of IMA or ADR for two months to generate derivative cells with mild, intermediate and severe resistance to the drugs as defined by their increasing resistance index (RI). PulseDIA-based quantitative proteomics was then employed to reveal the proteome changes in these resistant cells. In total, 7,082 proteotypic proteins from 98,232 peptides were identified and quantified from the dataset using four DIA software tools including OpenSWATH, Spectronaut, DIA-NN, and EncyclopeDIA. Sirtuin Signaling Pathway was found to be significantly enriched in both ADR- and IMA-resistant K562 cells. In particular, IDH2 was identified as a potential drug target correlated with the drug resistance phenotype, and its inhibition by the antagonist AGI-6780 reversed the acquired resistance in K562 cells to either ADR or IMA. Together, our study has implicated IDH2 as a potential target that can be therapeutically leveraged to alleviate the drug resistance in K562 cells when treated with IMA and ADR.

## Introduction

The treatment of chronic myeloid leukemia (CML) patients includes targeted therapy (tyrosine kinase inhibitors, TKIs), chemotherapy, biological therapy, hematopoietic cell transplant (HCT), and donor lymphocyte infusion (DLI) [1–3]. Chemotherapy inhibits the rapidly proliferating tumor cells by interfering with cell replication. However, drug resistance leads to failed chemotherapy treatment in 90% patients [4]. Adriamycin (ADR) is a traditional chemotherapeutic drug that disturbs the DNA replication process, which can be therapeutically leveraged against certain hematologic tumors. Using a K562 cell model, various mechanisms have been uncovered to explain ADR-induced drug resistance, such as transporter-mediated drug efflux [5, 6], altered mitochondrial function [7, 8], and changes in survival-related signaling pathway including EGFR/ERK/ NF-κB /PTEN/AKT [9–11].

As one of the most effective clinical regimens in the chronic phase, TKIs have dramatically improved survival rate in CML patients [12]. Indeed, imatinib (IMA) is among the first generation of TKIs and the first-line drug for CML treatment, which targets BCR-ABL1 and inhibits tumor growth [13]. Notably, the five-year survival rate of CML patients with IMA treatment was increased to 89% [14]. However, about 20-25% of CML patients showed a suboptimal response to IMA, who have likely developed drug resistance [15]. Several mechanisms have been proposed to explain the failed IMA treatment in CML patients, including for example altered conformation of the BCR-ABL1 kinase domain by mutations that reduces its binding affinity to IMA [12]. Other resistance mechanisms independent of BCR-ABL1 have also been reported, including P-glycoprotein upregulation, activation of alternative PI3K/AKT, JAK-2, or MAPK-signaling [16, 17], changes in the intracellular environment such as endoplasmic reticulum stress induced autophagy [18].

Drug resistance remains a clinical hurdle to traditional chemotherapy and targeted therapy [19]. Given its complex nature, both genetic mutations and non-genetic changes (such as epigenetics) may contribute to a drug-resistance phenotype [20]. To establish a cell model system and study drug resistance mechanism, we treated K562 cells with ADR or IMA and obtained a series of K562 cells with different drug sensitivities, which were used to recapitulate the process of acquired resistance.

The highly dynamic proteome, including changes in protein expression levels, protein-protein interactions and post-translational modifications, contributes to the phenotype in a cell [21]. Monitoring protein abundance through drug treatment may help reveal the underlying resistance mechanism [22, 23]. Mass spectrometry (MS)-based proteomics enables identification and quantification of thousands of proteins and thus offers unique insights into the dysregulated pathways [24]. In particularly, data-independent acquisition (DIA) is an effective proteomic method with rigorous quantitative accuracy and reproducibility [25, 26]. Based on DIA-MS, our group has recently developed PulseDIA-MS as an improved approach that utilizes gas phase fractionation to achieve a greater proteome depth [27].

In this work, we employed the pressure cycling technology (PCT)-based peptide preparation [28, 29], and the PulseDIA-MS strategy to quantify the dynamic proteome changes upon drug treatment, using time-series K562 drug-resistant cell line models treated by ADR and IMA, respectively. Due to the lack of correspondence information between precursor ions and fragment ions in DIA data, DIA data analysis has become a huge challenge. The different DIA tools have different scoring methods and core algorithm for peptides prediction, which may lead to certain technical deviations in the analytical results of the same DIA data [30]. To reduce the technical deviations of the quantitative proteome by DIA software tools, we quantified the proteome with four commonly used independent tools, Spectronaut [31], DIA-NN [32], EncyclopeDIA [33] and OpenSWATH [34]. With this cell model and quantitative approaches, we were able to pinpoint and characterize pathways that were significantly altered upon ADR or IMA treatment. Notably, we identified IDH2 a previously unknown potential target that can be therapeutically leveraged to reverse drug resistance in K562 cells.

## Results

### Establishment of drug-resistance cell models

Drug-resistant K562 cell models were developed as shown in the workflow (Fig. 1A). The parental K562 cells were sensitive to ADR and IMA treatment, and then treated with increasing concentrations of each drug for one, two, and four weeks to obtain derivative cell lines with differential drug sensitivities. Each model contains three time points during the drug resistance acquisition, with each time point containing cells showing a different degree of drug sensitivity (Fig. 1A, method).

**Figure 1.**
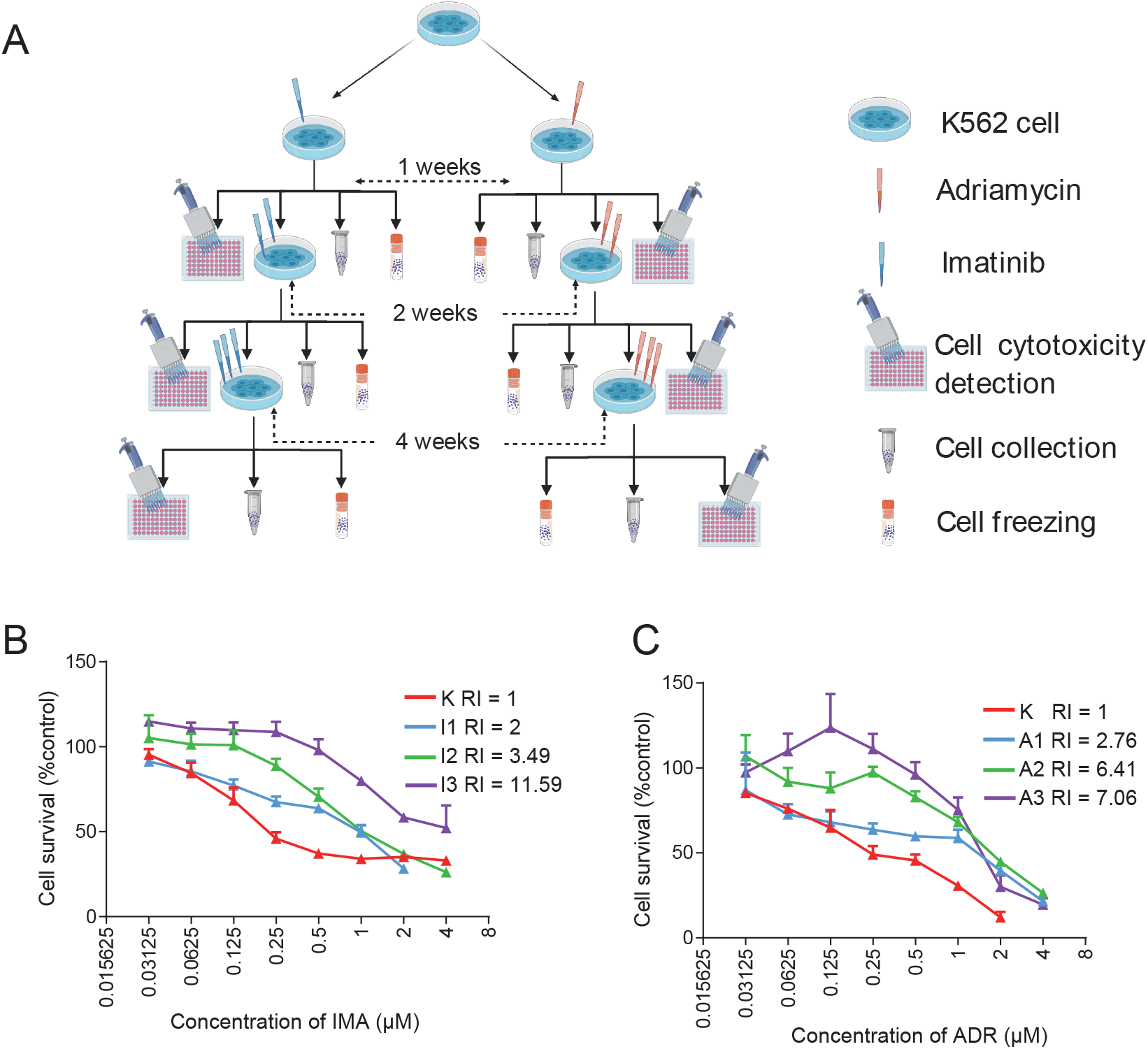
Establish derivitative K562 cells with mild, intermediate and severe resistance to ADR and IMA. (A) Overview of the drug resistance model. (B) Native K562 cells and IMA-resistant K562 cells from each of the three phases were treated with a series of cytotoxicity concentrations (4, 2, 1, 0.5……0 μM) IMA for 48 h. (C) Native K562 cells and ADR-resistant K562 cells from each of the three phases were treated with a series of concentrations (4, 2, 1, 0.5……0 μM) ADR for 48 h. The cell survival rate was calculated and plotted in each group. The downward shift of the survival curves by increasing treating concentration (I1, I2, I3 and A1, A2, A3) indicated suppressed proliferation.

At this point, we have generated derivative K562 cells with different degrees of IMA resistance: mild, intermediate and severe, which were defined by a resistant index (RI) at 2.76, 6.41 and 7.06 (Fig. 1B), respectively, as shown in Table 1. Similarly, we have mild, intermediate, and severe ADR-resistant K562 cells with a RI at 2.00, 3.49 and 11.59, respectively (Fig. 1C; Table 1). These K562 cells with different degrees of IMA and ADR resistance were all collected for PCT processing and further PulseDIA-MS analysis (Fig. 2A).

**Figure 2.**
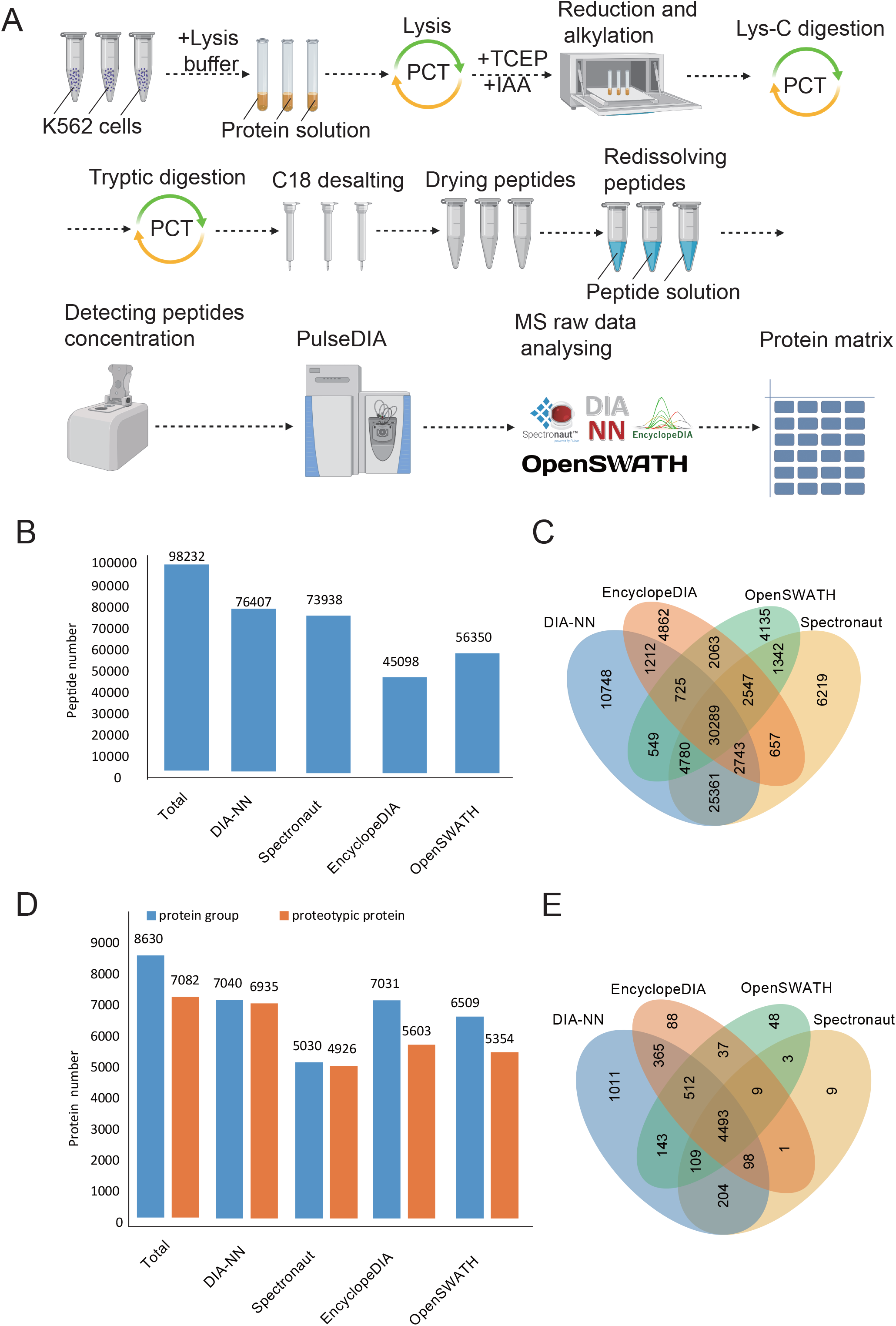
PulseDIA raw data acquisition and analysis. (A) The workflow of peptide extraction from samples and analysis of mass spectrometry raw data. (B) The numbers of identified peptides by four DIA software tools, and their Venn diagram (C). (D)The numbers of identified proteins by four DIA software, and their Venn diagram (E).

**Table 1.** IC50 values and resistance index of the derivative K562 cells.

| Model | IC50 (μM, 95%CI)<sup>a</sup> |  | Resistance Index<sup>b</sup> |
| --- | --- | --- | --- |
|  | IMA | ADR |  |
| Native K562 cells | 0.37(0.26 to 0.54) | 0.29 (0.24 to 0.35) | 1 |
| Model IMA phase 1 | 0.74 (0.61 to 0.90) | / | 2 |
| Model IMA phase 2 | 1.29 (1.06 to 1.58) | / | 3.49 |
| Model IMA phase 3 | 4.29 (3.03 to 6.36) | / | 11.59 |
| Model ADR phase 1 | / | 0.80 (0.51 to 1.24) | 2.76 |
| Model ADR phase 2 | / | 1.86 (1.5 to 2.30) | 6.41 |
| Model ADR phase 3 | / | 2.05 (1.32 to 3.32) | 7.06 |

<sup>a</sup>IC50 values represent the best-fit values and 95% CI of three independent experiments performed. <sup>b</sup>Resistance Index was calculated by dividing the best-fit IC50 values of native K562 cells in response to ADR and IMA to the best-fit IC50 values of model cells.

### Peptide and protein identification

PCT-assisted peptide preparation and PulseDIA-MS were then carried out to analyze the parental K562 cells and the derivative resistant cells in biological triplicates (Fig. 2A). In total, 98,232 peptides, 8630 protein groups,7082 proteotypic proteins were quantified from 26 independent PulseDIA-MS runs (Fig. 2B, D; Table. S1). 30,289 peptides and 4,493 proteotypic proteins were consistently quantified by four DIA software (Fig. 2C, E). We used these commonly identified proteins for the subsequent quantitative analysis of proteome changes during the development of drug resistance in K562 cells.

### Quality control (QC) of PulseDIA proteome dataset

We examined the reproducibility of PulseDIA data by calculating the Pearson correlation coefficient between technical duplicates and the coefficient of variation (CV) of the protein intensity among the biological triplicates. Using four different DIA analysis tools as described above, the five pairs of technical duplicates showed a strong correlation (r>0.9) (Fig. 3A), and the median CV of protein intensity among biological triplicates was around 20% (Fig. 3B). These results confirmed the high reproducibility of MS data acquired by the PulseDIA method. To compare the quantified result of the same peptides and proteins in the four DIA software, we calculated the Spearman correlation of the overlapped 30,289 peptides and 4,493 proteins in four DIA software quantified. At the peptide and protein levels, DIA-NN and Spectronaut exhibited the strongest correlation (r > 0.9), with the correlation of any two DIA software as no less than 0.85 (Fig. 3C, D). To check whether the quantified proteome thus acquired classify different drug resistance model, we performed also principal component analysis (PCA) analysis between the cells in ADR- and IMA-resistant cells, and parental K562 cells. The result showed that the two resistant cell lines and parental K562 cells were separated into three clusters as analyzed by the four software tools (Fig. 3E-H).

**Figure 3.**
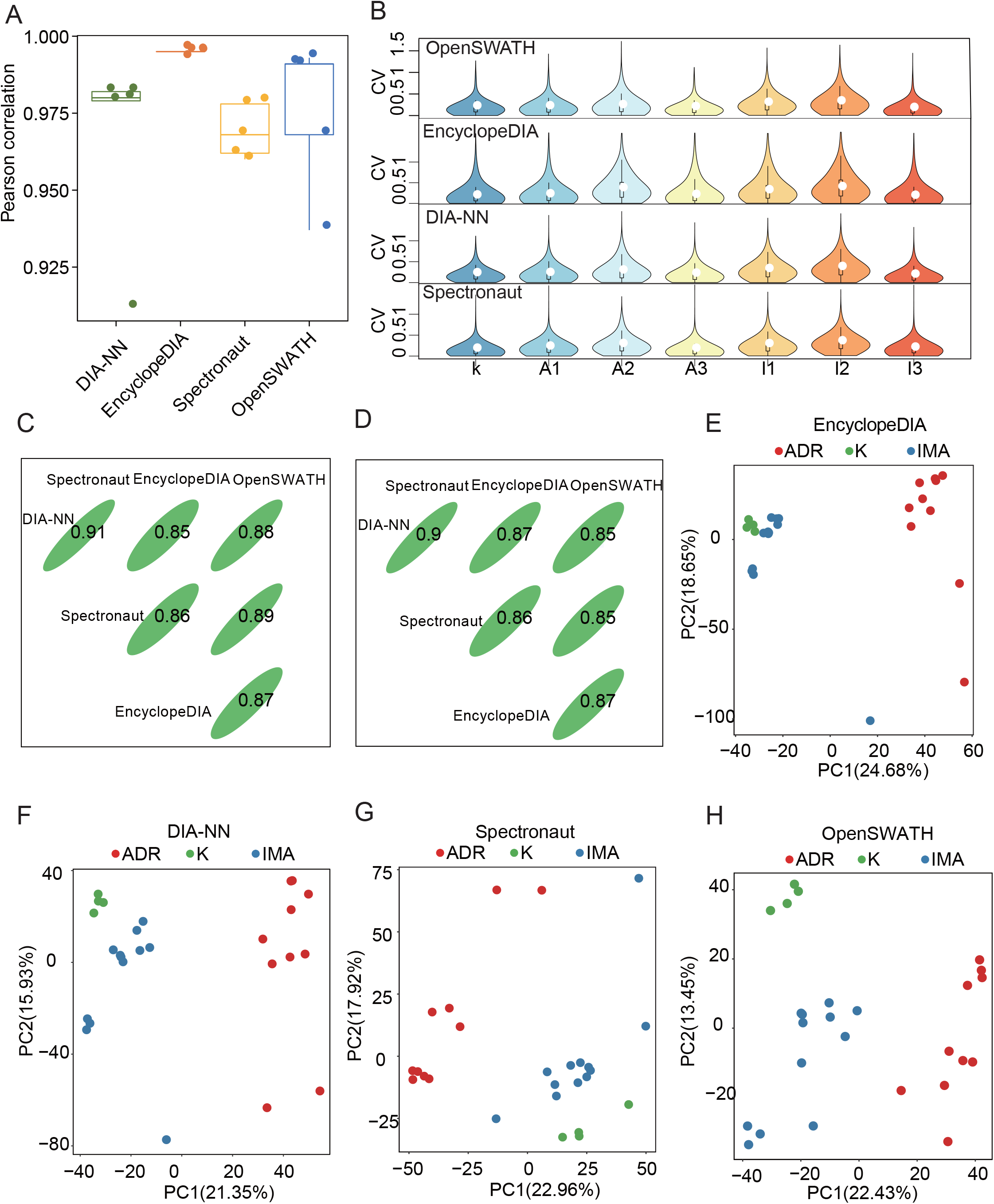
PulseDIA proteome data quality control (QC) analysis. (A)The box plot showed the Pearson correlation coefficient of 5 technical replicates in four DIA software quantification. (B) The violin plot showed the distribution of the CV of each protein’s quantitative values in three biological replicates. Three lines of the black box inside the violin represented lower quartile, median, higher quartile. (C) Spearman correlation coefficient of overlapped 30,289 peptides quantitative values in four DIA software. (D) Spearman correlation coefficient of overlapped 4493 proteotypic proteins quantitative values in four DIA software. (E, F, G, H) PCA plot shows the distribution of the sample in the first two principal component levels. The red, blue, green dots mean samples from model ADR, model IMA, native K562 cells, respectively.

### Dynamic proteomic changes during acquisition of drug resistance

To minimize the statistical variation for each analytic step, these commonly identified 4,493 proteins were selected and those with less than 25% missing ratio in each derivative resistant cell line were subject to the analysis of variance (ANOVA). Proteins with a p value less than 0.05 were selected for further downstream analysis.

By fuzzy c-means clustering [35, 36], we identified four clusters related to the resistance to ADR and IMA (Fig. S1; Fig. 4), amongst which only those that were continuously upregulated and downregulated were further selected and studied (Fig. 4). In addition, we selected 1035 (by Spectronaut), 1273 (by EncyclopeDIA), 1088 (by OpenSWATH), and 2161 (by DIA-NN) proteins related to resistance to ADR, (Table S2; Fig. 4A), and 1662 (EncyclopeDIA), 747 (OpenSWATH), 950 (Spectronaut) and 1027 (DIA-NN) proteins related to resistance to IMA (Table S3; Fig. 4B).

**Figure 4.**
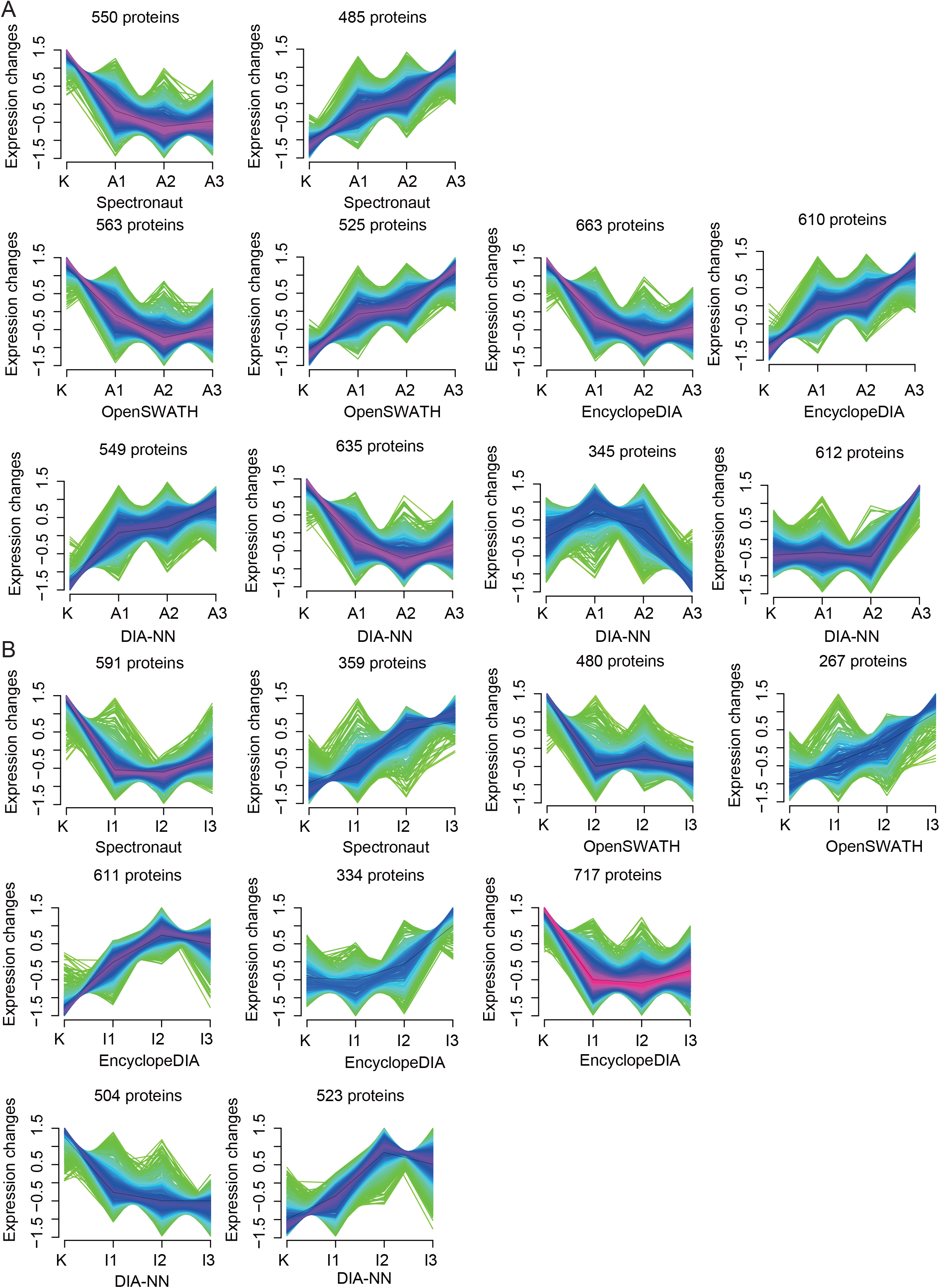
Protein cluster analysis. The cluster for proteins that were continuously upregulated and downregulated from K562 cells resistant to ADR (A) and IMA (B).The horizontal axis represented the progress of the model (K-native K562 cells, A1/I1-first phase of model ADR/IMA, A2/I2-second phase of model ADR/IMA, A3/I3-third phase of model ADR/IMA). The vertical axis represents proteins expression changes in each cluster.

### Activated sirtuin signaling pathway and abnormal mitochondrial function in drug resistant K562 cell

The selected proteins from the four DIA analytic tools as described above in resistance to ADR and IMA were further analyzed by IPA (Table S4, S5), with the top three pathways (sorted by p value) as listed in Figure 5A and 5B. Notably, EIF2 signaling and sirtuin signaling pathway were enriched from as least three DIA tools related to ADR resistance. In contrast, sirtuin signaling pathway and oxidative phosphorylation were frequently enriched related to IMA resitance (Fig. 5A, B). These data indicate that the sirtuin signaling pathway was significantly enriched in both types of drug resistance with a positive activation mode (Z-score>0, Table 2).

**Figure 5.**
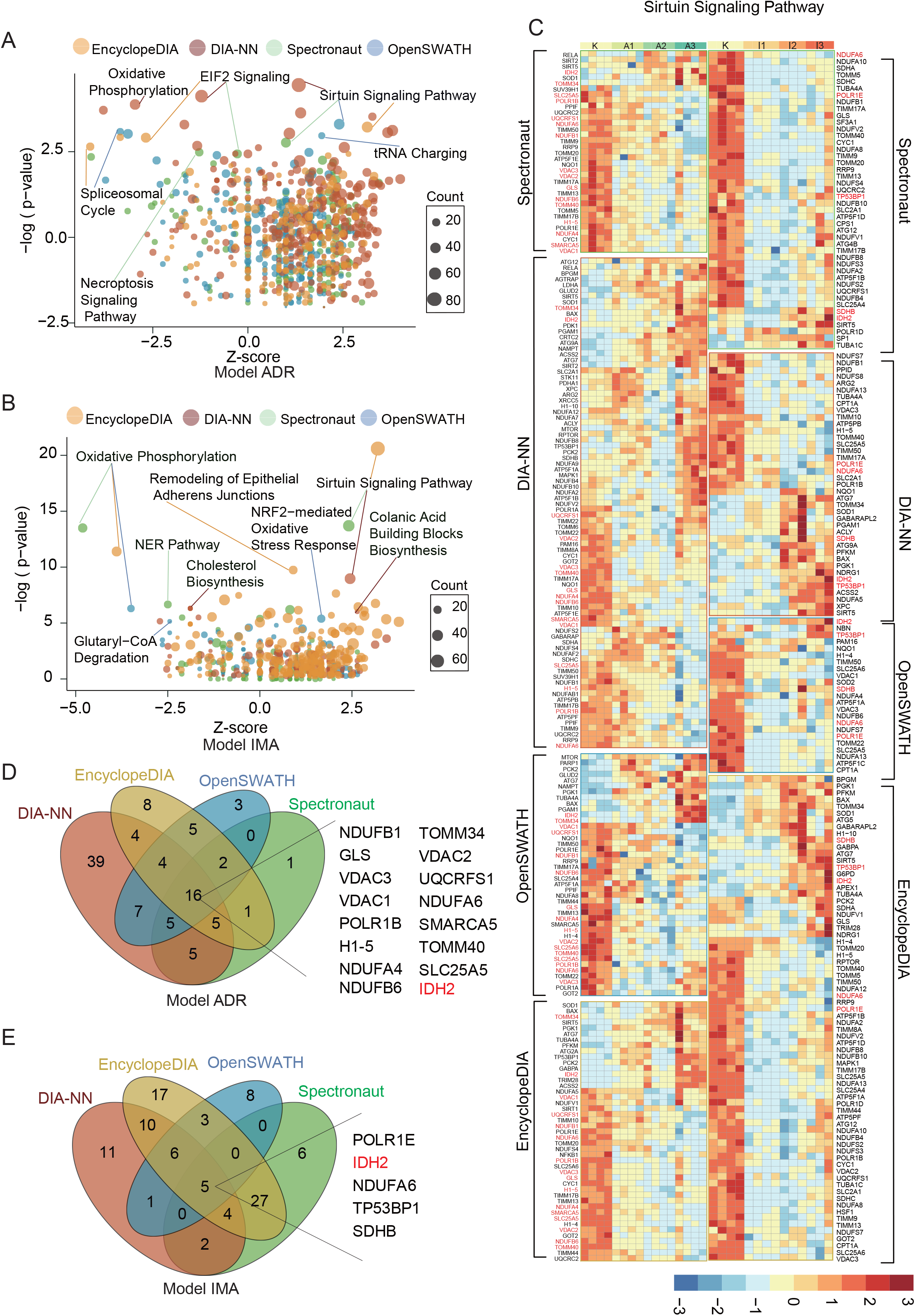
Enriched proteins that exhibit continuous changes as identified by IPA in ADR- and IMA-resistant cells. (A) In ADR-resistant cells, the most significantly changed 3 pathways were enriched by IPA from four DIA software. (B) In IMA-resistant cells, the most significantly changed 3 pathways were enriched by IPA from four DIA software. (C) The heatmap showed the proteins (p value<0.05 calculated by ANOVA) involved in Sirtuin signaling pathway from four DIA software expression in IMA- and ADR-resistant cells, with the overlapped quantified proteins being highlighted in red. Each row indicated a protein, and each column indicated a sample. The protein intensity matrix was normalized by Z-score and colored in the heatmap. (D) Venn diagram showed the 16 overlapped proteins involved in Sirtuin Signaling Pathway in ADR-resistant cells. (E) Venn diagram showed the 5 overlapped proteins involved in Sirtuin Signaling Pathway in IMA-resistant cells.

**Table 2.** The enrichment statistics of Sirtuin Signaling Pathway in four DIA software.

| Software | P value<sup>a</sup> | Ratio<sup>b</sup> | Z-score<sup>c</sup> | Model |
| --- | --- | --- | --- | --- |
| OpenSWATH | 4.48 | 0.0825 | 1.155 | Model IMA |
| Spectronaut | 13.7 | 0.155 | 2.414 | Model IMA |
| DIA-NN | 8.97 | 0.134 | 2.449 | Model IMA |
| EncyclopeDIA | 20.6 | 0.251 | 3.202 | Model IMA |
| OpenSWATH | 9.87 | 0.144 | 2.4 | Model ADR |
| Spectronaut | 6.8 | 0.12 | 1 | Model ADR |
| DIA-NN | 21.8 | 0.296 | 1.336 | Model ADR |
| EncyclopeDIA | 9.31 | 0.155 | 3.138 | Model ADR |

<sup>a</sup>P-value is the result of Fisher's exact test. <sup>b</sup>Ratio is the number of proteins from different DIA software that maps to the pathway divided by the total number of proteins that map to the same pathway. <sup>c</sup>Z-score is calculated by the IPA z-score algorithm to predict the direction of change for the function. An absolute z-score of $\geq 2$ is considered significant.

We then focused on the proteins involved in the sirtuin signaling pathway, whose expression levels were shown in a heatmap (Fig. 5C). As a result, 16 proteins, which were involeved in sirtuin signaling pathway, were quantified between all four DIA analytic tools in model ADR (Fig. 5D), and 5 proteins in model IMA (Fig. 5E). The overlapped proteins between all four DIA analytic tools showed the same expression pattern in the analysis results of four DIA software (Fig. 5C). Among the overlapped proteins, 13 out 16 related to ADR resistance were involved in maintaining mitochondrial function, and three out of five related to IMA resistance participated in the formation of the mitochondrial respiratory chain (Fig. 6A).

**Figure 6.**
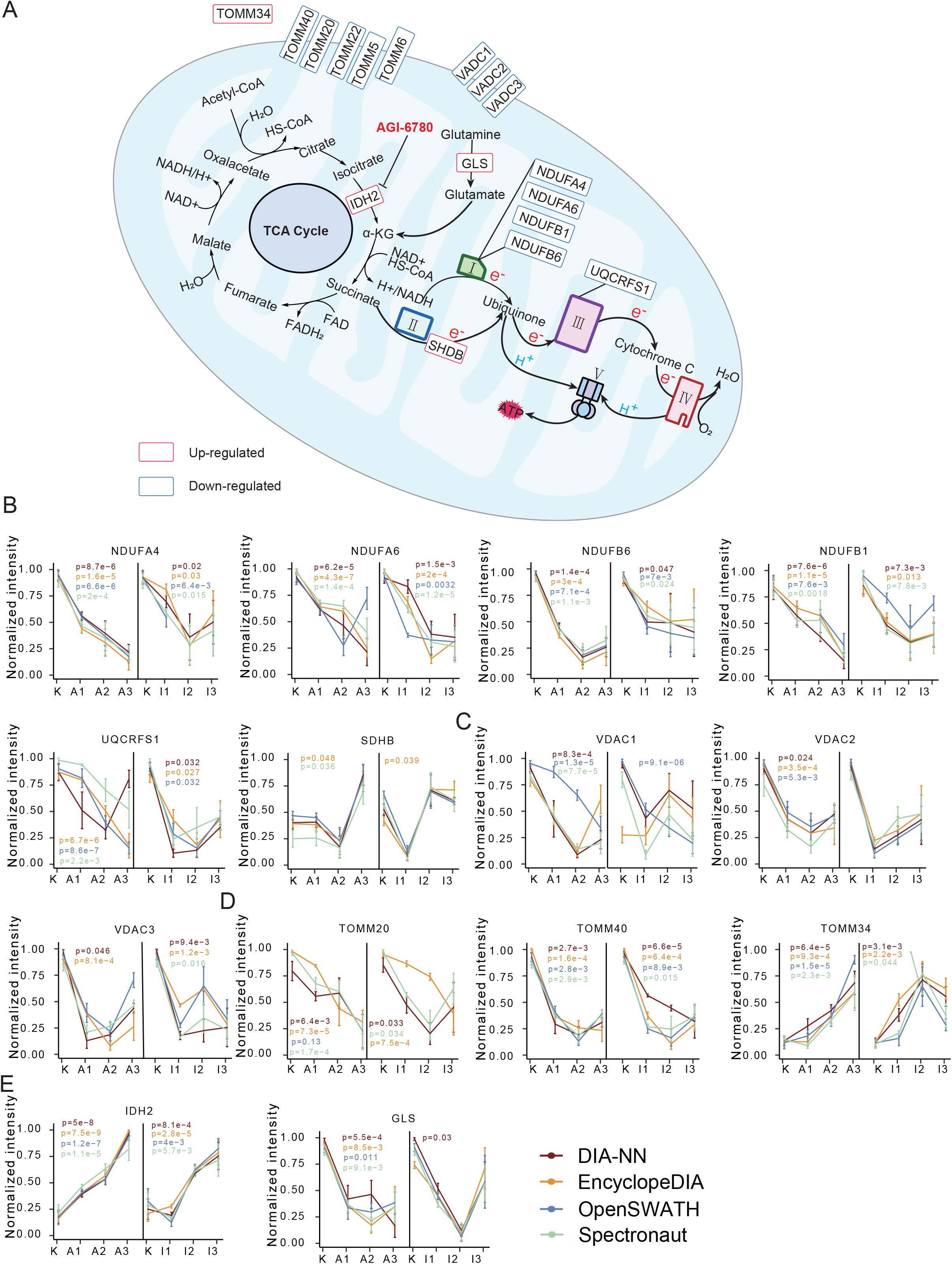
Proteins involevd in mitochondrial function. (A) Schemathic diagram of TCA cycle and electron respiratory chain in mitochondria. (B) Normalized protein expression in ADR- and IMA-resistant cells from four DIA software.

NDUFB1, NDUFB6, NDUFA4, and NDUFA6 are the subunits of NADH-ubiquinone oxidoreductase (Complex I) of the mitochondrial respiratory chain (Fig. 6A) [37]. UQCRFS1 is a subunit of ubiquinol-cytochrome c oxidoreductase (complex III) (Fig. 6A). The main physiological function of Complex I and Complex III is to oxidize NADH, transfer electrons to ubiquinone and from ubiquinone to cytochrome C, and carry out the next electron transfer in the mitochondrial respiratory chain (Fig. 5A) [38]. In addition, Complex 1and Complex III also regulate mitochondria production of reactive oxygen species (ROS) [39]. Our results showed decreased expression levels of these Complex I and III components (Fig. 6A, B), suggesting low oxidative stress levels in drug-resistant cells.

We also identified upregulation of succinate dehydrogenase complex iron sulfur subunit B (SDHB), which participates in the electron transfer process of succinate dehydrogenase (Complex II) from succinate to ubiquinone (Fig. 6A, B), and when mutated, is closely related to pheochromocytoma [40].

Voltage dependent anion channel proteins (VDACs) are the most abundant protein [41] on the outer mitochondrial membrane (Fig. 6A), and function to maintain free diffusion of small molecules inside and outside the mitochondrial membrane [42]. In tumor cells, the interaction between VDAC and hexokinase inhibits apoptosis [43], and therefore targeting both molecules could potentially offer improved anti-tumor benefits. Our results showed that the three VDAC isoforms, namely VDAC1, VDAC2, and VDAC3, were significantly downregulated upon acquision of ADR resistance (Fig. 6C), suggesting that targeting VDAC may not be an effective strategy in drug-resistant tumor cells.

The translocase of the outer mitochondrial membrane complex proteins (TOMMs) regulates entry of mitochondrial protein precursors into the mitochondria cytoplasm (Fig. 6A) [44]. Our data showed that the subunits of TOMMs, TOMM40, TOMM5, TOMM6, TOMM22, and TOMM20, were significantly downregulated in cells resistant to ADR or IMA (Fig. 5C; Fig. 6A, D). In contrast, we found increased expression of TOMM34 upon acquision of drug resistance (Fig. 5D). Located in the cytoplasm, TOMM34 is known to interact with HSP70 and HSP90, and regulate the activity of ATPase [45]. High expression of TOMM34 has been found in colon cancer[46], breast cancer[47], ovarian cancer [48].

Glutaminase (GLS) can convert glutamine into glutamic acid, which constitutes the major source for α-ketoglutarate (α-KG) production(Fig. 6A) [49, 50], which promotes cell differentiation through dioxygenases [51]. Our results showed significantly down-regulated expression of GLS when K562 cells became resistant to ADR (Fig. 6E).

Isocitrate Dehydrogenase (NADP(+)) 2 (IDH2) catalyzes the oxidative decarboxylation of isocitrate into α-ketoglutarate (α-KG) in tricarboxylic acid cycle (TCA cycle), and NADPH was synchronously produced at the biochemical process [52] (Fig. 6A). NADPH is essential in protecting cells from oxidative damage[53]. We found the IDH2 abundance increased upon resistance to both ADR and IMA (Fig. 6E), indicating that IDH2 overexpression may promote cellular resistance against high doses of both therapeutic drugs.

Taken together, our results showed significantly changed abundance of proteins involved in mitochondria function that likely mediate survival of drug-resistant cells, which may be therapeutically targeted to enhance drug sensitivity and response in tumor cells.

### IDH2 is a potential target for reversing drug resistance in K562 cells

As discussed above, among the dysregulated proteins, IDH2 was upregulated in K562 cells resistant to both ADR and IMA. To validate its biological function, we utilized a selective inhibitor of IDH2 [54], AGI-6780, and treated sensitive or resistant K562 cells with ADR and IMA alone or combined with AGI-6780, followed by monitoring cell survival with a cytotoxicity assay. Our results showed that, compared with ADR and IMA treatment alone, combination with AGI-6780 did not affect the survival in the parental sensitive K562 cells (Fig. 7A, B; Table 3). However, in resistant K562 cells, the sensitivity to ADR and IMA was significantly increased (IMA+AGI-6780, IC_50_=0.53μM; ADR+AGI-6780, IC_50_=0.29μM), when combined with AGI-6780. We further show in IMA-resistant cells the reversal index of 2 μM and 4 μM AGI-6780 was 1.92 and 2.94, respectively (Fig. 7C; Table 3), whereas in ADR-resistant cells, the reversal index of 2 μM and 4 μM AGI-6780 was 1.60 and 2.74, respectively (Fig. 7D; Table 3). Taken together, the cytotoxicity assay confirmed the ability of AGI-6780 to reverse drug resistance *in vitro*, implicating IDH2 as a potential target to improve drug sensitibity and clinical responses in patients.

**Figure 7.**
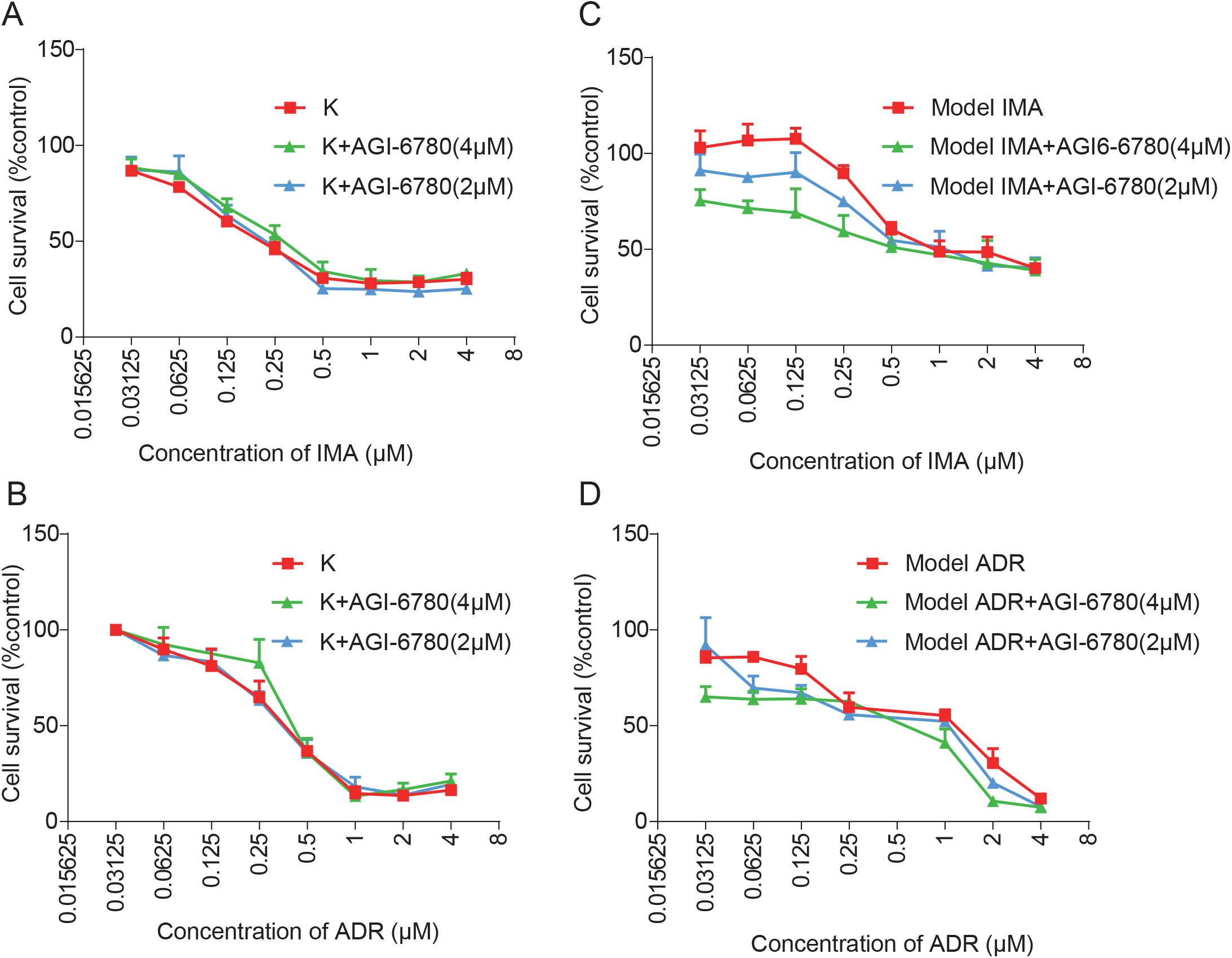
IDH2 enhanced the sensitivity to ADR and IMA in K562 cells. (A) and (C) The parental sensitive and IMA-resisant K562 cells were treated with various concentrations (4, 2, 1, 0.5……0 μM) IMA alone or combined with 4 μM or 2 μM AGI-6780 for 48h. (B) and (D) The parental sensitive and ADR-resisant K562 cells were treated with various concentrations (4, 2, 1, 0.5……0 μM) ADR alone or combined with 4 μM or 2 μM AGI-6780 for 48h. The cell survival rate was calculated and plotted in each group. The downward shift of the survival curves indicated suppressed proliferation. Columns are expressed as mean ± SD.

**Table 3.**
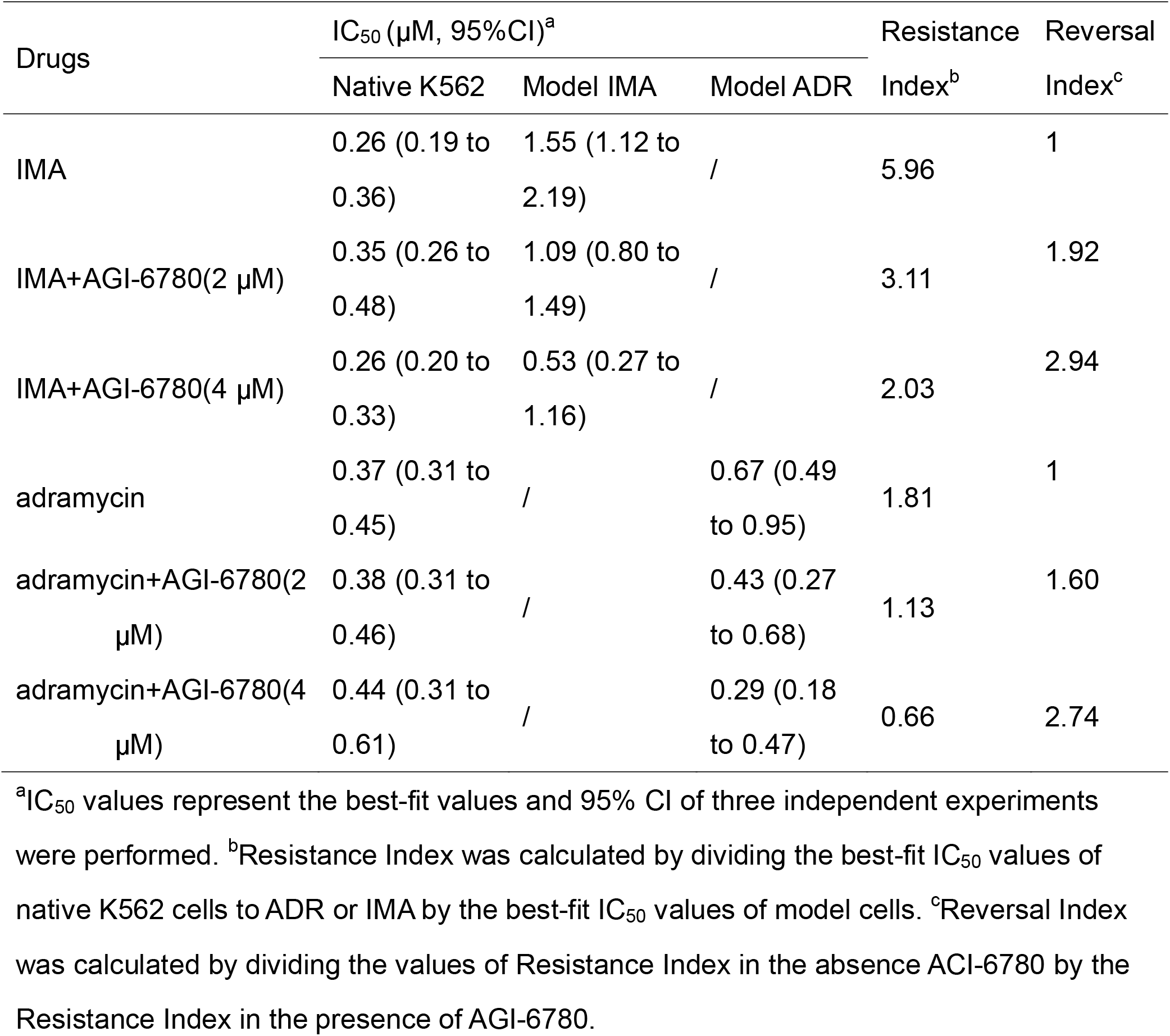
IC_50_ values and reverse index of the derivative resistant K562 cells.

| Drugs | IC<sub>50</sub> (μM, 95%CI)<sup>a</sup> |  |  | Resistance | Reversal |
| --- | --- | --- | --- | --- | --- |
|  | Native K562 | Model IMA | Model ADR | Index<sup>b</sup> | Index<sup>c</sup> |
| IMA | 0.26 (0.19 to 0.36) | 1.55 (1.12 to 2.19) | / | 5.96 | 1 |
| IMA+AGI-6780(2 μM) | 0.35 (0.26 to 0.48) | 1.09 (0.80 to 1.49) | / | 3.11 | 1.92 |
| IMA+AGI-6780(4 μM) | 0.26 (0.20 to 0.33) | 0.53 (0.27 to 1.16) | / | 2.03 | 2.94 |
| adramycin | 0.37 (0.31 to 0.45) | / | 0.67 (0.49 to 0.95) | 1.81 | 1 |
| adramycin+AGI-6780(2 μM) | 0.38 (0.31 to 0.46) | / | 0.43 (0.27 to 0.68) | 1.13 | 1.60 |
| adramycin+AGI-6780(4 μM) | 0.44 (0.31 to 0.61) | / | 0.29 (0.18 to 0.47) | 0.66 | 2.74 |

<sup>a</sup>IC<sub>50</sub> values represent the best-fit values and 95% CI of three independent experiments were performed. <sup>b</sup>Resistance Index was calculated by dividing the best-fit IC<sub>50</sub> values of native K562 cells to ADR or IMA by the best-fit IC<sub>50</sub> values of model cells. <sup>c</sup>Reversal Index was calculated by dividing the values of Resistance Index in the absence AGI-6780 by the Resistance Index in the presence of AGI-6780.

## Discussion

Two major paradigms have been widely proposed to elucidate the underlying mechanism of drug resistance. One is based on the principle of Darwinian selection, which posits that a fraction of tumor cells is inherently more tolerant and thus become enriched after drug treatment, thereby gradually developing and exhibiting a drug-resistant phenotype. The other mechanism, which is called Lamarckian induction, proposes that drug treatment induces a resistance phenotype in tumor cells [55]. In our resistant cell line model, our focused was on the proteins that showed continuous and consistent changes upon acquisition drug resistance. The underlying cause for these changes is not immediately clear from our work, but our results demonstrated that altered expression levels of these proteins were responsible for the drug resistance phenotype in K562 cells, which could uncover potentially important targets for managing and reversing drug resistance in clinics.

While establishing the resistance cell lines, we also constantly monitored drug sensitivity in the parental sensitive K562 cells (Fig. 1B, C). Different from other similar studies [56], we did not consider morphology changes as a parameter, primarily because K562 cells are suspension cells in the medium in a spherical shape while the drug resistant phenotype was established. After K562 cells with mild, intermediate and severe resistance to ADR and IMA were developed and collected, we performed PCT-based peptides extraction and PulseDIA-based mass spectrometry raw data acquisition. The efficient and prompt sample preparation under high pressure (lysis in 45K psi; digest in 20K psi) by PCT ensures that the process of peptides extraction is reproducible and stable [29].

Spectronaut[31], DIA-NN[32], EncyclopeDIA[33], and OpenSWATH[34] utilize different algorithms to analyze the DIA data, and the results from these distinct analytic tools are therefore expected to be different. To avoid the algorithm bias from an individual software analysis, we used all four tools to analyze our PulseDIA data. Our results showed that the coefficient of variation of protein intensity in three biological replicates was around 20% (Fig. 3B). For PulseDIA based mass spectrometry raw data acquisition, we identified and quantified 83.1% (7082/8524) proteotypic proteins of the DDA library from 26 samples by the four DIA datasets, and 63.4% (4493/7082) proteotypic proteins were detected by all four tools (Table. S1; Fig. 2D, E). The PulseDIA data stability was assessed by the Pearson correlation of two technical replicates, which was no less than 0.9 (Fig. 3A). The PCA result demonstrated that the resistant K562 cells can be discriminated from the parental cells at the whole proteome level (Fig. 2E–H). These high-quality mass spectrometry data provides a key basis for our subsequent data mining analysis.

We independently analyzed 4493 proteins identified by all four tools. Based on cluster analysis and ANOVA (p≤0.05), we selected proteins that exhibited continuous changes during development of IMA and ADR resistance (Table S2, S3; Fig. 4), followed by IPA analysis. Pathway analysis revealed common characteristics associated with drug resistance. Significant changed stress signaling pathways including Oxidative Phosphorylation and the Sirtuin Signaling Pathway (Fig. 5A, B) are known to enhance the resistance and plasticity of tumor cells[57–59]. We then decided to focus on the identified proteins involved in the Sirtuin Signaling Pathway as identified by all four DIA tools, including IDH2, NDUFB1, NDUFB6, NDUFA4, NDUFA6, SHDB, and GLS, which are involved in mitochondrial ATP generation (Fig. 5D, E; Fig. 6A)[60, 61]. These results suggest that inhibiting ATP production and blocking P-gp energy supply could potentially enable reversal of drug resistance [5, 62].

IDH2 mutation catalyzes D-2-hydroxyglutarate (2-HG) production, which leads to competitive inhibition of α-KG-dependent DNA demethylases and consequently promotes tumorigenesis [51]. Preclinical experiments have shown that IDH2 inhibitors promote leukemic cell differentiation in 40% of the patients with relapsed/refractory AML[54]. Here, we used the selective IDH2 inhibitor AGI-6780, and demonstrated IDH2 as a potential target for reversing drug resistance in tumor cells. Recent research has shown that 5 μM AGI-6780 selectively impaired wild type IDH2 enzymatic activity in multiple myeloma cells in vitro[61]. In contrast, our data showed no effects on cell proliferation even 48 h after 4 μM AGI-6780 treatment in either sensitive or resistant K562 cells (Fig. S2). As a potential therapeutic avenue for reversing drug resistance, IDH2 inhibitors are specifically efficacious in resistant tumors cells.

Derivative cell lines that have developed drug resistance could provide an important and informative model to help elucidate the underlying mechanism that can be strategically leveraged to manipulate and improve the sensitivity of tumor cells to therapeutic treatment. Single-cell proteomics studies of melanoma-derived cells with different levels of drug sensitivity revealed changes in intracellular signals before drug resistance has developed [22]. By constructing a cisplatin-resistant neuroblastoma cell line, Olga et al. characterized the epithelial to mesenchymal transition during the development of drug resistance [56]. Herein, we have developed multiple K562 cell lines with different degrees of resistance to ADR and IMA treatment, which could be used for preclinical investigation of the molecular and cellular mechanisms that underlie the therapeutic resistance in CML patients. We further identified and characterized IDH2 as a potentially useful target that can be utilized to enhance tumor cell sensitivity to targeted treatment.

## Acknowledgements

This work is supported by grants from National Key R&D Program of China (No. 2020YFE0202200), the National Natural Science Foundation of China (81874324, 81972492, 21904107), Zhejiang Provincial Natural Science Foundation for Distinguished Young Scholars (LR19C050001), Hangzhou Agriculture and Society Advancement Program (20190101A04), and Westlake Education Foundation. We thank Westlake University Supercomputer Center for assistance in data storage and computation.

## Data availability

All data are available in the manuscript or the supplementary material. The proteomics data are deposited in ProteomeXchange Consortium. All the data will be publicly released upon publication. The project data analysis codes are deposited in GitHub.

## Materials and Methods

### Establishment of drug-resistance K562 cell models

The parental sensitive K562 cells were purchased from Nanjing KeyGen Biotech Co., Ltd. (Nanjing, China) and authenticated via ShortTandem Repeat (STR) profiling by Shanghai Biowing Applied Biotechnology Co., Ltd (Shanghai, China) on Feb. 28, 2019. The parental K562 cells were cultured in RPMI medium (Cromwell, USA) with 10% fetal bovine serum (Waltham, USA) and 1% penicillin-streptomycin (HyClone, USA) at 37 °C with 5% CO2 and 95% humidity.

The building of drug-resistance K562 cell models includes three phases (Fig. 1A). In the first phase, 1×10^5^ K562 cells were treated with 0.1 μM ADR (A1) or IMA (I1). After one week, K562 cells were cultured to about 10 million cells. Then drugs were removed by centrifugation and cells were divided into four aliquots for cell cytotoxicity determination, subsequent drug-resistance induction, cell collection, and cell freezing. In the second phase, the concentration of ADR and IMA was increased to 0.4 μM (A2) and 0.8 μM (I2), respectively. The treatment lasted for two weeks. In the third phase, the concentration of ADR and IMA increased to 0.8 μM (A3) and 1.6 μM (I3), respectively. The treatment lasted for four weeks. In this way, we established two drug-resistance K562 models for ADR and IMA, separately.

### Cytotoxicity assay

The cytotoxicity of Adriamycin (CAS: 25316-40-9, meilunbio, Dalian, China), imatinib mesylate (CAS: 220127-57-1, meilunbio, Dalian, China) or AGI-6780 (CAS: 1432660-47-3, MedChemExpress, New Jersey, USA) to native K562 cells and the model cells were detected by cell counting kit-8 (CCK-8) (Bimake, Houston, USA). Cells were planted into 96-well plates in a density of 5000 cells/100uL medium/well. After 24h, cells were treated with drugs for 48h. After 48h, 10μL CCK-8 was added to each well and incubated about 2h in dark. The absorbance value of each well was determined by a Synergy™ H1 (BioTek, State of Vermont, USA) at 450nm. The IC50 value was calculated by SPSS 22.0.

### PCT based peptide extraction

The workflow of PCT based peptide extraction is described as Shao et al [29]. For each sample, 500,000 cells were harvested and cleaned by PBS three times to remove all traces of fetal bovine serum.

Cells were transferred into PCT-Microube (Pressure Biosciences Inc.) with 30uL lysis buffer, 5 uL 1ug/uL DNAase (STEMCELL) and 15uL 0.1M ammonium bicarbonate (ABB) (GENERAL-REAGENT^®^). The lysis buffer includes 6M urea (SIGMA-ALDRICH^®^) and 2 M thiourea (SIGMA-ALDRICH^®^. Then the cells were lysis by Barocycler NEP2320-45k (PressureBioSciences Inc.) with 90 cycles containing 25 s of 45,000 p.s.i. high pressure plus 10 s at ambient pressure, at 30 °C The lysate solution was added 2.5 μL 800 mM tris(2-carboxyethyl)phosphine (ALDRICH^®^) and 5 μL 100 mM iodoacetamide (SIGMA^®^) to in PCT-Micro tube to dilute into a final concentration of 10 mM and 40 mM, followed by a 30 min incubation in the dark with gentle vortexing (800 rpm) at room temperature in shaker.

After reduction and alkylation, the proteins solution were added with 57.5μL 0.1 M ABB and 25 μL 0.1 mg/mL Lys-C (Hualishi) and digested using Barocycler NEP2320-45k (Pressure Biosciences Inc., MA, USA) with 45 cycles containing 50 s of 20,000 p.s.i. high pressure plus 10 s at ambient pressure, at 30 °C.

After Lys-C digestion, the solution was added with 10 μL 0.2 mg/mL trypsin (Hualishi) and were tryptic digested by Barocycler NEP2320-45k with 90 cycles containing 50 s of 20,000 p.s.i. high pressure plus 10 s at ambient pressure, at 30. Then, 15 μL 10% trifluoroacetic acid (Fisher Scientific®) was added to the lysate solution at a final concentration of 1% to stop digestion.

Then the digested peptides were cleaned in micro spin columns (The Nest Group Inc.) and dried in CentriVap DNA Vacuum Concentrators (LABCONCO). Peptides were redissolved in ms buffer (0.1% formic acid and 2% acetonitrile in HPLC water). The peptide concentration was measured using ScanDrop^2^ (Analytik Jena).

### PulseDIA mass spectrometry

The PulseDIA mass spectrometry method was performed as previously described [63]. The redissolved peptides of each sample were acquired and analyzed by EASY-nLC™ 1200 System (Thermo Fisher Scientific™) coupled to a QE HF-X mass spectrometer (Thermo Fisher Scientific™). The MS1 was acquired in an m/z range of 390 to 1210 with the resolution at 60,000, AGC target of 3e6, and the maximum ion injection time of 80 ms. The MS2 was performed with the resolution at 30,000, AGC target of 1e6, and the maximum ion injection time of 50 ms. Different from the conventional DIA method (the 400-1200 m/z mass range is divided into 24 windows) [64], the PulseDIA symmetrically divided each window into four parts and acquired the data of each part independently. For one sample, we acquired four injections with different MS2 range. For each pulse acquisition, 0.2 μg of peptides was injected and separated across a linear 45 min LC gradient (from 8% to 40% buffer B) at a flowrate of 300 nL/min (precolumn, 3μm, 100 Å, 20mm*75μm i.d.; analytical column, 1.9um, 120 Å,150mm*75um i.d.). Buffer A was HPLC-grade water containing 0.1% FA, and buffer B was 80%ACN, 20%H2O containing 0.1%FA.

### Quality control samples

Cells with mild, intermediate and severe resistance to ADR and IMA were analyzed in 3 duplicates for peptide extraction and acquisition as biological replicates. Five samples were repeatedly injected by PulseDIA as the technical replicates to evaluate the data quality.

### Generating Spectral library for DIA-MS

To build the spectral library, we acquired 37 data-dependent acquisition (DDA) files including 10 in-solution digestion files, 7 PCT-assisted digestion files and 20 high-PH fraction files on a QE-HFX mass spectrometer in DDA mode. Library was built by Spectronaut (version: 13.5.190902.43655) for Spectronaut and DIA-NN analysis. Library was built by OpenSWATH (version 2.0) for OpenSWATH and EncyclopeDIA analysis.

For Spectronaut library building, Carbamidomethyl (C) was set as the fixed modification, and Acetyl (Protein N-term) and Oxidation (M) were set as the variable modification. Data were searched against the SwissProt Human database in February 2018. Q value (FDR) cutoff on precursor and protein was 0.01. Finally, a K562 library including identified 191,008 precursors, 133,025 peptides, 8226 protein groups and 8313 proteins was built.

For OpenSWATH library building, the pFind[65] (version 2.8) was used as a search engine with the parameters including carbamidomethyl (C) as a fixed modification and methionine (M) and oxidation (O) as a variable modification. Other parameters were performed as default. Data were searched against the SwissProt Human database in February 2018 and further filtered the data with FDR□≤□1%. DDA library was built according to the workflow [66] (http://openswath.org/en/latest/docs/pqp.html#id3). Finally, a K562 library including identified 110,583 transition groups, 84,548 target peptides, 84,910 decoy peptides, 9511 target protein groups, 9575 decoy protein groups and 7935 target proteotypic proteins was built.

Base on the analysis of K562 Spectral library, we finally identified a total of 8524 proteotypic proteins and up to 10,732 protein groups.

### PulseDIA data analysis

Library-based PulseDIA data analysis was performed by Spectronaut, OpenSWATH, DIA-NN, and EncyclopeDIA.

For Spectronaut (version: 13.5.190902.43655) analysis, the default setting of library-based DIA analysis was used for PulseDIA analysis. PulseDIA analysis was performed according to the standard workflow in Spectronaut (Manual for Spectronaut, available on the Biognosys website). The FDR was set to 1% at the peptide precursor level.

For DIA-NN analysis, we used the same library with Spectronaut analysis. Library search was performed according to the DIA-NN manual (https://github.com/vdemichev/DiaNN/blob/master/DIA-NN%20GUI%20manual.pdf). The precursor FDR was set as 0.01.

For OpenSWATH (version: 2.4) analysis, the retention time extraction window was 300 seconds. Peptide precursors that were identified by OpenSWATH and pyprophet with qvalue=0.01.

For EncyclopeDIA (versions: 0.9.0) analysis, the library was converted from OpenSWATH library. The precursor, fragment, and library mass tolerance were set as 10 ppm for the PulseDIA data.

The peptide matrixes from four DIA software tools were converted to protein matrixed by R code (https://github.com/Allen188/PulseDIA/blob/master/Pulsedia_DIANNresult_combine.R)

### Bioinformatics analysis

Soft cluster analysis was performed by R/Bioconductor package Mfuzz [36]. The average protein intensity of the parental K562 cells and each type of resistant cells as the input data for clustering. The time-series were separated according to the cell sensitivity to ADR or IMA, with the initial being the parental K562 cells. Ingenuity Pathway Analysis (IPA, QIAGEN) was performed to outline the significant canonical pathways [67].

### Calculation of IC_50_ values

The IC_50_ values were calculated by a non-linear least squares regression model to fit the data log (inhibitor) vs. normalized response in GraphPad Prism 6. The statistical significance in different types of resistant cells was determined by one-way analysis of variance (ANOVA), and the p value of <0.05 was regarded as statistically significant. In IPA, the p-value was calculated using the right-tailed Fisher’s exact test and a P value less than 0.05 is considered significant[68].

## Legends of Supplementary Files

**Supplementary Figure 1.**
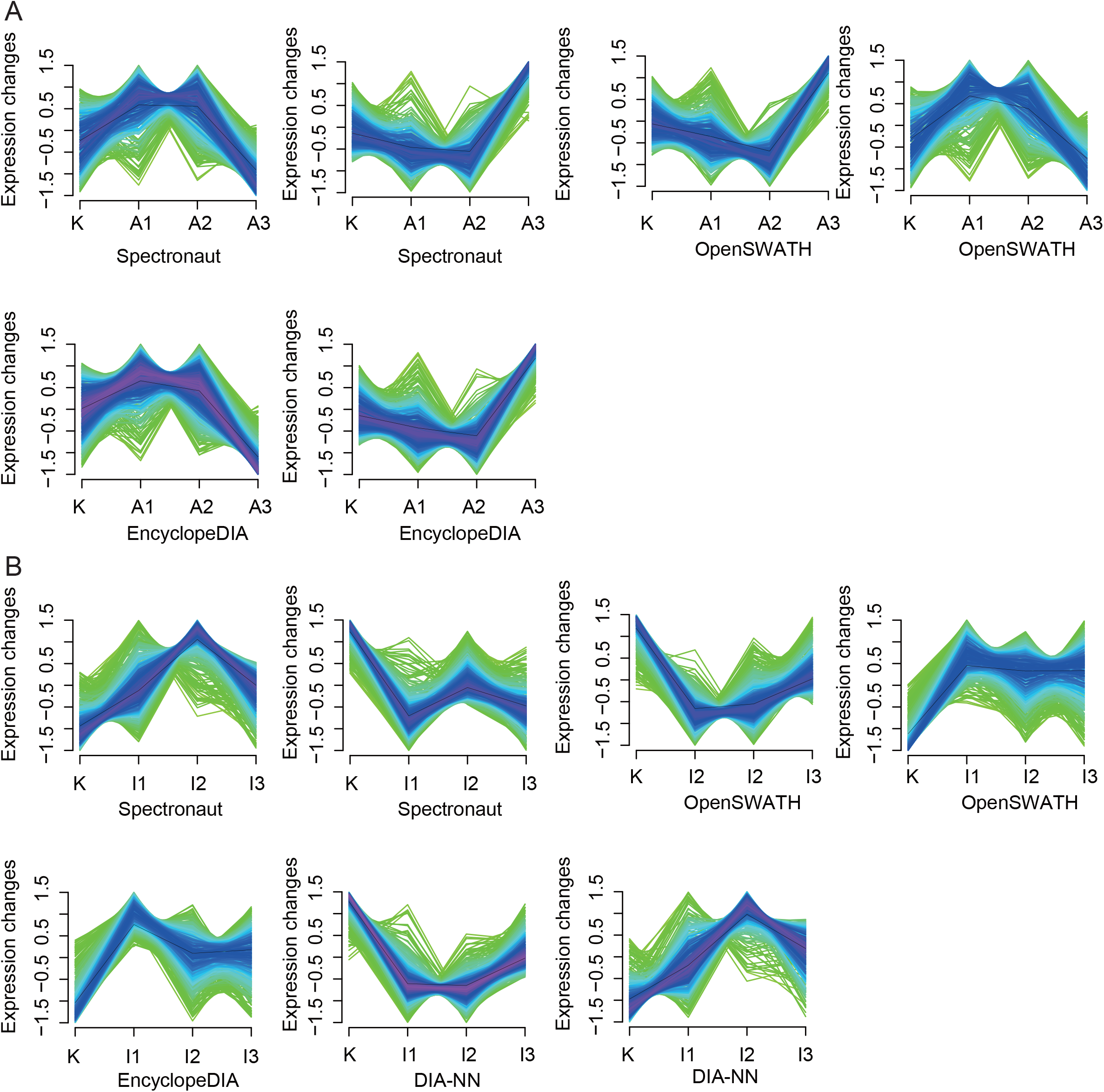
Protein cluster analysis. The protein cluster from model ADR (A) and IMA (B).The horizontal axis represented the progress of the model (K-native K562 cells, A1/I1-first phase of model ADR/IMA, A2/I2-second phase of model ADR/IMA, A3/I3-third phase of model ADR/IMA). The vertical axis represented changes in proteins expression in each cluster.

**Supplementary Figure 2.**
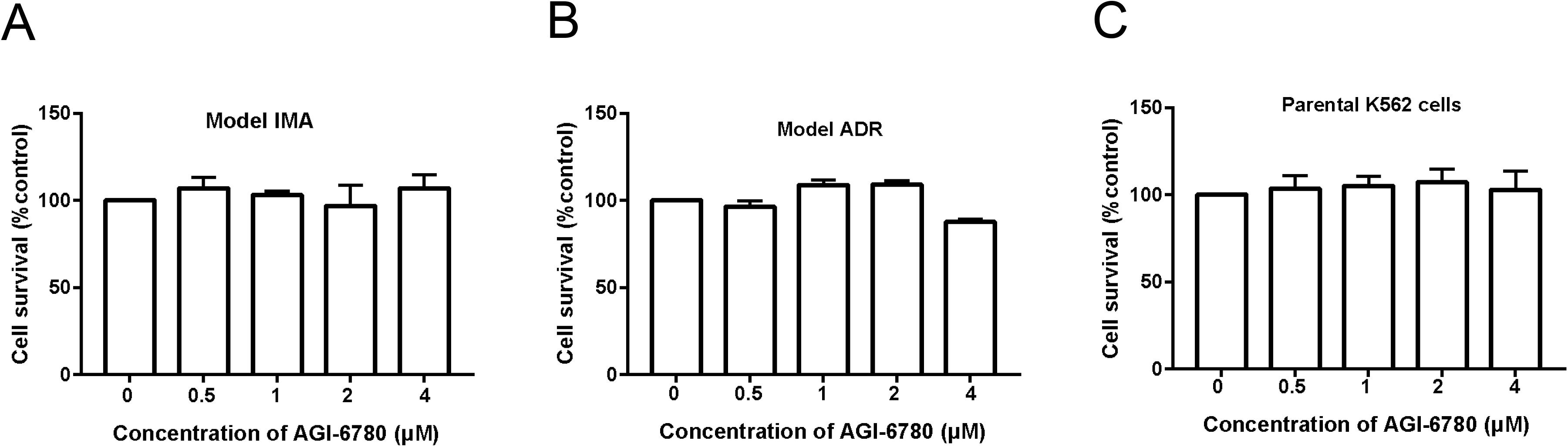
Changes in cell survival after different concentrations AGI-6780 (0, 0.5, 1, 2 4 μM) treated IMA-resistant (A), ADR-resistant (B) and parental K562 cell (C) for 48h.

**Supplementary Table 1.** Protein matrix of quantified proteins in 26 samples by DIA-NN, Spectronaut, EncyclopeDIA and OpenSWATH, the quantitative protein abundance was log2 converted.

**Supplementary Table 2.** In ADR-resistant cells, the list of continuously changed proteins (p < 0.05) with associated p values.

**Supplementary Table 3.** In IMA-resistant cells, the list of continuously changed proteins (p < 0.05) with associated p values.

**Supplementary Table. 4** IPA analysis of continuously changed proteins from ADR-resistant cells.

**Supplementary Table. 5** IPA analysis of continuously changed proteins from IMA-resistant cells.

